# Schizophrenia, variability, and the Anna Karenina principle

**DOI:** 10.1101/2025.09.30.679570

**Authors:** Michael Murphy, Dost Öngür

**Affiliations:** McLean Hospital; Belmont, MA, 02447, USA; Department of Psychiatry, Harvard Medical School; Boston, MA 02115

**Author notes:** Corresponding author: Michael Murphy, 115 Mill St, Belmont MA 02447. Dr. Murphy reports no financial relationships with commercial interests. Dr Öngür has recieved personal fees from Neumora Inc, Guggenheim LLC, Boehringer Ingelheim, and Rapport Therapeutics. This work was funded by National Institutes of Health grant K23MH118565 (MM). We thank Mark Halko for his helpful comments on this manuscript. MM designed the study, analyzed the data, and wrote the study. DO wrote the study.

## Abstract

In neuroimaging studies of people with schizophrenia there is often higher within group variance in the patient group compared to the control group. This is counterintuitive – why would a subset of people selected because they all have the same disease be more varied than the general population? We used simulated data and real neuroimaging data to identify a potential cause of elevated variance in populations of patients with schizophrenia. We demonstrated that elevated variance can arise within variables that are unrelated to disease status simply because people with a set of neurological perturbations that cause schizophrenia are more likely to have higher numbers of perturbations overall. Additionally, we showed that observed elevated variances in people with schizophrenia can be reproduced by models that only rely on perturbation count.

These results highlight an important barrier in our attempts to understand the pathophysiology of schizophrenia. Standard statistical practices in schizophrenia research do not account for the fact that schizophrenia is, at every level of analysis that has been studied, highly heterogeneous. This heterogeneity by itself is sufficient to produce elevated variances. Our work suggests that the most effective way to prevent schizophrenia may not be to identify and mitigate specific pathologies but rather to reduce the impact of broadly damaging factors such as those associated with poverty.

## Introduction

Neuroimaging studies of schizophrenia often report that many measures have a higher intra-group variability in the case group compared to control group. For example, Kozhemiako et al. (2024) reported that, after controlling for clinical, cognitive, and medication factors, 645 out of 4746 sleep electroencephalography variables had a higher variance in the schizophrenia group while only 44 variables had a higher variance in the control group (*1*). Similar findings have been reported in functional magnetic resonance connectivity studies (*2*). These elevated variances are counterintuitive. Why would a subpopulation that was selected to share a set of common features (i.e., disease status) have higher variances than the population as a whole? One possibility is that neuroimaging methods may cleave schizophrenia into discrete sub-populations. For example, one group of patients with schizophrenia may have increased theta power and decreased alpha power compared to controls while another group of patients has decreased theta and increased alpha power. In both groups, there are abnormal theta and alpha power values.

Statistical testing may not show mean differences in these power bands between cases and controls but will likely show elevated variance in the case group compared to the control group. If this were the case, schizophrenia subgroups could potentially be derived from the data.

However, studies to date have not reported the presence of clearly defined schizophrenia subgroups, rather, EEG variables in schizophrenia are distributed on a spectrum (*1*). Large scale neuroimaging studies have used clustering algorithms like k-means to partition schizophrenia into a small number of putative classes (*3*). These analyses have so far produced classes that poorly capture the variance within the schizophrenia population and do not map onto clusters in other measures (*4*).

Alternatively, it may be that increased variance in patient populations is driven by differences in exposure to medication. Psychiatric medication can impact neuroimaging measures in a variety of ways with different medications having different effects (*5*). Furthermore, two patients who are prescribed the same dose of the same medication may have different adherence which introduces an additional source of variability. Studies that have controlled for medication exposure still show elevated variance in cases compared to controls. Finally, elevated variance may arise from different compensatory responses to the pathophysiology that drives schizophrenia. It may be that the brain utilizes a large set of compensatory responses and that these responses, while not directly related to schizophrenia, are nonetheless a result of it. All of these explanations for increased variance propose that the specific parameters with elevated variance tell us something about the pathophysiology of schizophrenia. However, there is yet another possibility – the elevated variances reflect something about the analysis space and not about the individual parameters. Schizophrenia is binary at exactly one level, the level of case-control status. Individuals are either diagnosed with schizophrenia or they are not. At every other level of analysis, the term “schizophrenia” refers to a set of multiple distinct multi-hit phenomena. The multiplicity of schizophrenias has important implications for how we should interpret variance at a given level of analysis. We should apply the Anna Karenina principle from systems analysis – that at many levels of analysis there are so many different schizophrenias that there may not be useful common features between them. At these levels of analysis, the variance in the case population is likely to be higher than the variance in the control population. This elevated variance arises from 1) the absence of common features across cases and 2) the fact that the more perturbations a person experiences, the more likely they are to experience a set of perturbations that leads to schizophrenia.

## Materials and Methods

Modeled and real data were analyzed using Matlab (Mathworks, Natick MA, USA). Models were coded as described in the text. Details of EEG data collection, preprocessing, and sample demographics can be found in (10). Details of fMRI data collection, preprocessing, and sample demographics can be found in (11).

### Data Analysis

For both simulated and real data, we used Bartlett’s test to compare the variances within variables between case and control groups. For comparisons between groups, we used unpaired t-tests. We tested for normality of hit count distributions with the Shapiro-Wilk test. We used the Kolmogorov-Smirnov test to determine of modeled hit count distributions agreed with empirically derived ones.

## Results

### Models suggest that people with schizophrenia have more non-schizophrenia symptoms than healthy controls

Schizophrenia is defined clinically. If case-control status is the binary “top” layer, then clinical symptoms are the next layer down. A simplified version of the DSM-5 criteria for schizophrenia is that a person needs to have two or more of the following symptoms: hallucinations, delusions, disorganized speech, disorganized behavior, and negative symptoms with at least one of either hallucinations, delusions, or disorganized speech (*6*). Already, we encounter multiple different schizophrenias. A person with schizophrenia may have disorganized speech, disorganized behavior, and negative symptoms. A different person with schizophrenia may have hallucinations and delusions. Not only do these individuals have different sets of symptoms, but they have no symptoms in common and yet both have schizophrenia. At the level of symptoms, there are 25 different combinations of symptoms that meet the criteria for schizophrenia. In this simplified version of the DSM, these 25 combinations capture the entirety of schizophrenia – there is no unexplained signal in the patient population. This feature is unique to this level of analysis. That is, as of now, there is no set of neuroimaging, genetic, plasma or other markers that perfectly captures all schizophrenias.

To model this, we can create a binary vector for each individual where each entry represents a symptom and the number 1 means it is present and the number 0 means it is absent. The DSM contains 628 unique symptoms and therefore each vector has 628 entries with the first five entries corresponding to hallucinations, delusions, disorganized speech, disorganized behavior, and negative symptoms respectively (*8*). Thus, our individual with disorganized speech, disorganized behavior, and negative symptoms can be represented as [0 0 1 1 1…] and the person with hallucinations and delusions as [1 1 0 0 0…]. Note that in this simplified example, the values of entries after the first five play no role in determining whether a person has schizophrenia. In the following discussion, if a symptom is present and there is a 1 in the vector, we will refer to it as a “hit”. The number of hits in an individual is the hit count for that person.

We can then use this simple frame to construct a model population. Each modeled individual is given some number of symptoms each of which are chosen from a set of probability distributions. For the first symptom assigned to an individual, we can use data about the frequency of each symptom in the population to create a weighted distribution to draw from.

Once a symptom is selected, we can use data about the co-occurrence of symptoms (for example, in one large study, 66% of people who reported hallucinations also reported having delusions while 15% of people with delusions also had hallucinations) (*7*). Given that there are no large-scale studies of symptom co-occurrence that include every unique symptom in the DSM, we can model this more generally by creating for each symptom a subset of symptoms with which it is tightly affiliated above a background level of chance affiliations. This model of symptom co-occurrence is broadly consistent with network models of psychopathology (*9*).

### The distribution of hits drives between-group differences in variance

There are multiple parameter choices in such a model including the shape of the distribution of symptom counts and the shapes of the symptom-specific symptom likelihoods. We will model the hit counts as a normal distribution. For the associations between symptoms, we construct a probability vector for each symptom. The entries in this vector indicate the probability of having each other symptom if the current symptom is present. We construct this vector by creating a probability vector with each entry drawn from a normal distribution. We then randomly select a small, random number of entries to have much higher values than the average value of the vector. Finally, we renormalize the vector to sum to 1. This process creates a probability vector with a small number of highly correlated entries above a background of lower correlations. In this way, we model that some symptoms may be highly associated (as seen in (*7*)).

By adjusting the parameters of the normal distribution of hit counts, we can adjust the rate of schizophrenia in our modeled population. Our parameter choices impact whether cases or controls have more variables with higher variance and how strong this effect is. To illustrate this point, we can simulate multiple studies drawing from modeled populations of 100000 individuals. Consider two normal distributions for hits, each of which yields a schizophrenia prevalence of 0.51%, consistent with epidemiological studies. With one normal curve for hit distribution (mean = 14 symptoms/person, standard deviation = 9, skew = 0.6, kurtosis =3, Fig. 1a), we can use Bartlett’s test to measure variance differences between groups and find that, on average, for 30 studies each with sample size of 25 cases and 25 controls, 208 variables (modeled symptoms) have statistically significantly more variance in the case group while 109 variables have statistically significantly more variance in the control group. With a different curve (mean = 15 symptoms/person, standard deviation = 3, skew = 0, kurtosis = 3, Fig. 1b), we instead find that 145 variables have statistically significantly more variance in the case group while 159 have statistically significantly more variance in the control group. Note that in both simulations, cases have more hits on average than controls do. More dramatic effects can be seen if we model hit counts using other distributions – for example, a similar set of simulations drawn from an exponential distribution that produces the same prevalence (Fig. 1c) yields 315 variables with more variance in cases compared to 64 such variables in the controls. Clearly, the distribution of hits in a population critically determines how variance is distributed across the case and control subpopulations. Furthermore, knowledge of the differences in variance between case and control groups provides information about the distribution of hits in the overall population.

**Fig. 1.**
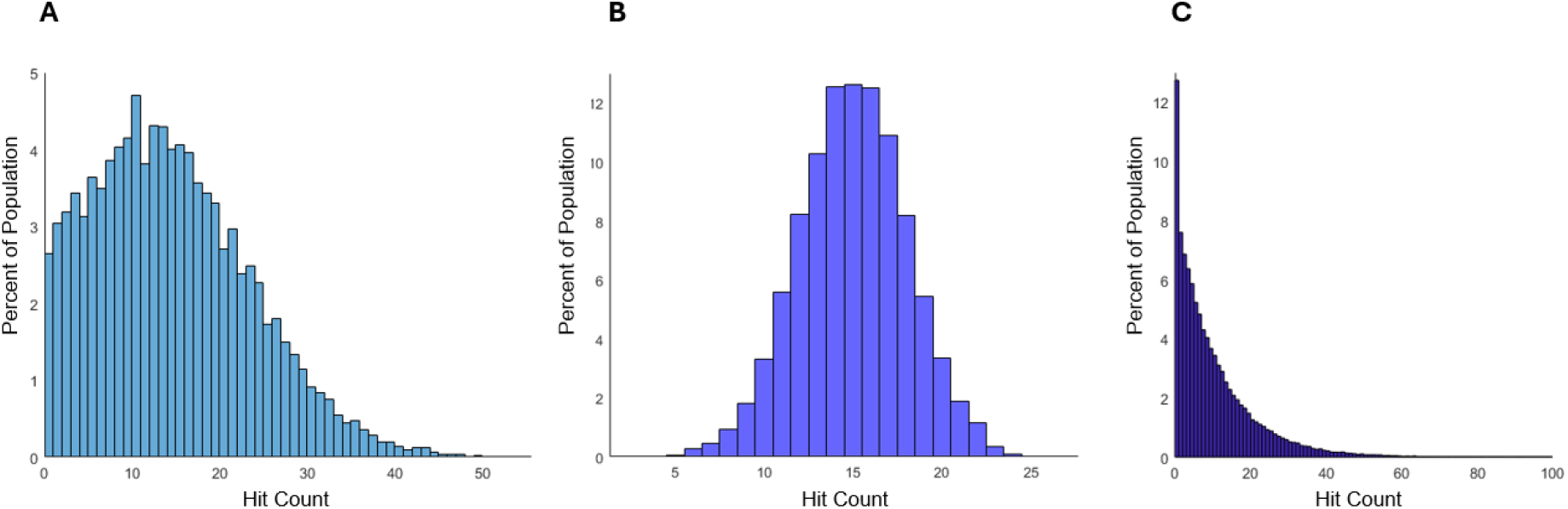
Different hit count distributions can produce similar disease prevalence but different distributions of variance between cases and controls. These three hit count distributions all produce a schizophrenia prevalence of 0.51% given the same set of correlations between symptoms. (**A)** and (**B):** Normal distributions with different means, standard deviations, skew, and kurtosis. (**C):** An exponential distribution.

### Differences in variance in neuroimaging data can be modeled using only hit count

How can we be sure that the variance differences in individual variables between cases and controls reflect something about schizophrenia specifically rather than reflecting that the schizophrenia population is more likely to have all sorts of perturbations and therefore higher variance? If the variance in the individual variables is important to schizophrenia, there is likely to be some meaning in the correlation structure of the variables (that is, the matrix of correlations between the EEG variables). In particular, if elevated variances in schizophrenia reflect schizophrenia subtypes or subtype-specific responses then we would expect there to be a complicated high-order correlation structure where the values of some variables are greatly impacted by specific sets of values in other variables. Variables that are related to some forms of schizophrenia are likely to have some correlation to a subset of other variables that are related to some of the same forms of schizophrenia. We directly interrogate this possibility using real neuroimaging data.

With real data, we can estimate some of the parameters of the hit space curve. In a recent study, we collected resting-state EEG data from 79 controls and 114 cases and calculated 1091 different EEG variables (10). Comparing the variance in cases and controls with Bartlett’s test showed that 190 variables had more variability in cases and 51 had more variability in controls. We can test the effect of hit number and correlation structure by simulating data. We binarize the data by creating a binary vector for each participant where each entry is an EEG variable and is scored a 1 (and called a hit) if it is at least 20% different from the subgroup mean. We then train a decision tree on this matrix to identify cases and controls. We then identify the mean hit number per person (188.6), standard deviation of the hits per person (55.5), skew (0.45), and kurtosis (2.7) of the distribution of hits per person in the control group (Fig. 2). We use these parameters to construct a normal distribution of hit counts (note that a Shapiro-Wilk test on the real hit count does not reject the null hypothesis that the data is normally distributed). To eliminate the effect of high-order correlations, we replace the correlation structure with random correlations. We simulate a population drawn from this distribution and simulate 30 studies with 79 controls and 114 cases. We find that on average there are 112.2 variables with higher variance in cases and 30.3 variables with higher variance in controls (p < 10^-26^ unpaired t-test). This ratio is comparable to the ratio from the real data suggesting that any correlations between variables have limited impact. Furthermore, these results suggest that some, but not all, of the elevated variance in the cases is related to the cases having more hits. Some of the remaining differences in variances between cases and controls are related to how the variables are correlated with each other in real data and some of this correlation structure may be related to various forms of schizophrenia.

**Fig. 2.**
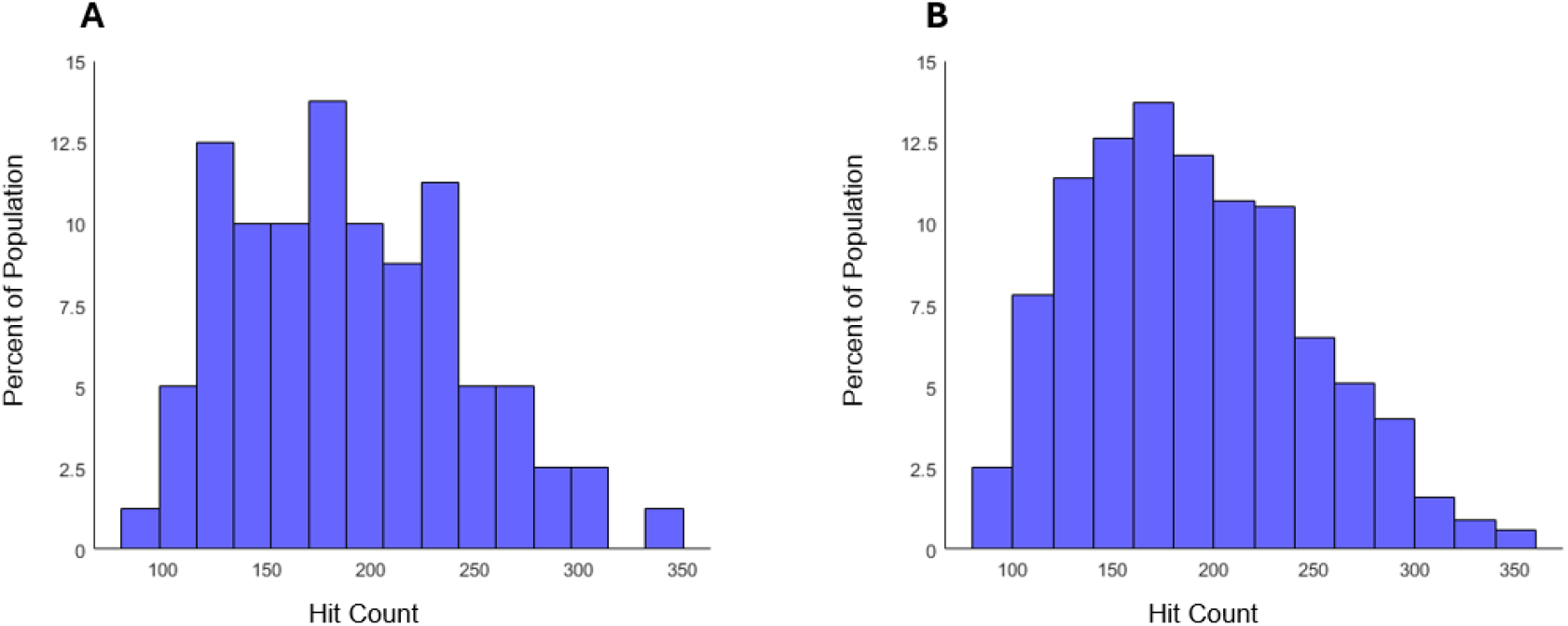
EEG data hit count distribution can be modeled with a normal distribution. (**A)** A histogram of hit counts from the control data. (**B)**: A histogram of hit counts from a population that is drawn from a normal curve fitted to the real data.

This effect is not unique to EEG. We analyzed 9730 resting-state fMRI connectivity variables from 92 subjects with schizophrenia spectrum disorders and 92 control participants (11) . 1142 variables had more statistically significantly more variance in cases compared to controls and 616 variables had more variance in controls. We binarized the data in a similar fashion to the EEG analysis described above and noted that the hit count distribution is not normally distributed (p<.05, Shapiro-Wilk test). Instead, we modeled the hit count distribution as a mix of Gaussians which adequately captured the structure of the distribution (p>.1, Kolmogorov-Smirnov test, Fig. 3). We used this data to simulate a population and again simulated 30 studies with 92 cases and 92 controls. When using a random correlation matrix between the variables in our simulated data, we find that, on average, 1350.0 variables have more variance in cases and 748.4 variables have more variance in controls (p < 10^-20^ unpaired t-test). Again, higher variances in cases may be at least partially explained by cases having more hits.

**Fig. 3.**
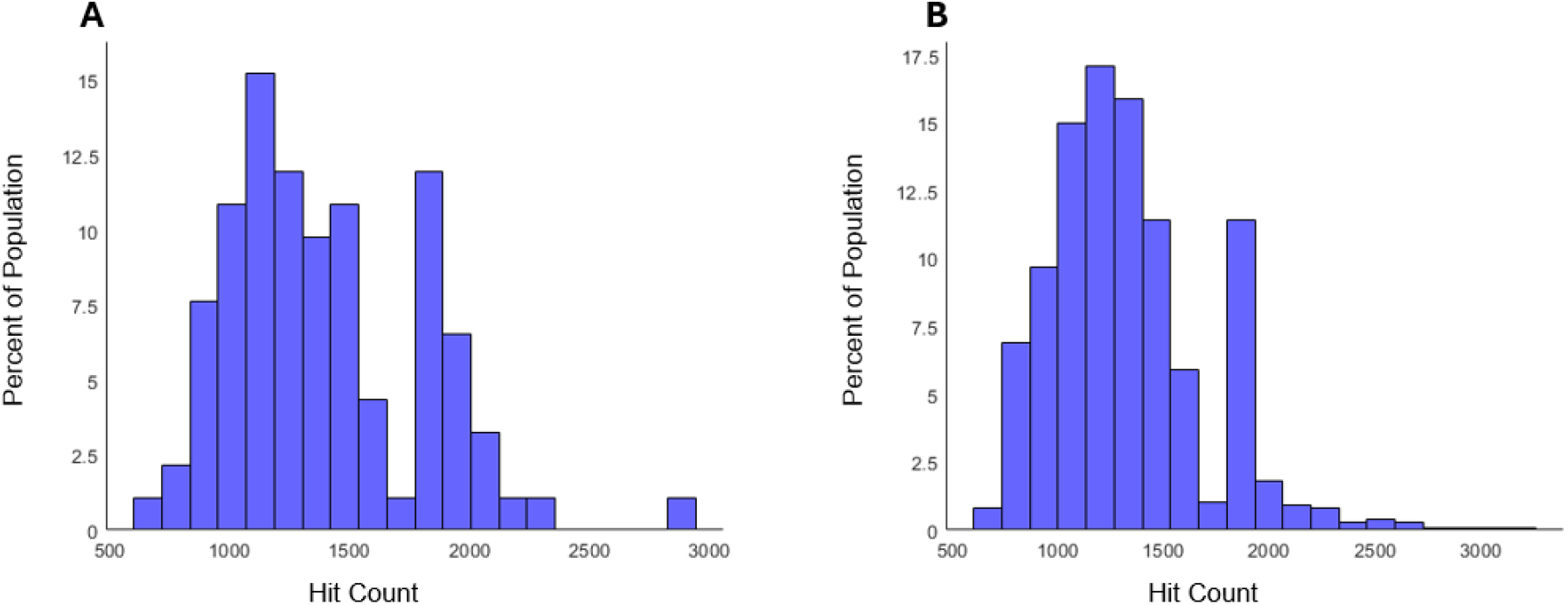
fMRI data hit count distribution cannot be modeled by a normal distribution, but can be modeled by a mix of normal distributions. **(A)** A histogram of hit counts from the control data. **(B)** The hit counts from a population that is drawn from a mix of normal distributions fitted to the real data.

## Discussion

These simulations do not disprove the possibility of schizophrenia-specific factors that increase variance in neuroimaging measures. For example, people with schizophrenia move more during MRI scans which can make these measurements noisier (12). However, higher variability has been observed across multiple scales and units of measurement in schizophrenia (for example, (13,14)). The highly linear relationship between per-chromosome single nucleotide polymorphism (SNP) contributions to schizophrenia and chromosome length is consistent with the possibility that some portion of the observed heritability is related to overall SNP count (15).

We have demonstrated that we can generate higher variances in the cases even if we have random correlations between variables. These simulation results indicate that we should acknowledge the possibility that at least some differences in variance between cases and controls are related to different amounts of hits in cases compared to controls. People who experience more hits are also more likely to encounter a sequence of hits that leads to a diagnosis of schizophrenia. Thinking about variance in this way leads to strategies for addressing complicated challenges. Despite decades of research, our understanding of the pathophysiology of schizophrenia remains inadequate and has not translated into meaningful interventions for people with schizophrenia. Implicit in our research programs is an assumption, or perhaps a hope, that there is a tractable number of schizophrenias. This assumption stands in opposition to the longstanding appreciation that schizophrenia is a heterogenous syndrome. Here, we present evidence that the assumption of a small number of schizophrenias is unsupported by our data.

Widely used statistical techniques do not and cannot account for a complex syndrome like schizophrenia. However, our work also supports the idea that it is not necessary to understand the etiologies of the syndrome to decrease its prevalence. Rather, we should focus on decreasing the hit counts in the general population. Interventions like ameliorating childhood poverty, decreasing substance use, and removing toxins from the environment will lead to lower levels of schizophrenia in the population. This will be true even if we think of these hits as causing essentially random damage that only occasionally produces schizophrenia. That is, factors that are broadly and indiscriminately damaging will be major drivers of schizophrenia simply by driving up hit counts in the population. Schizophrenia is not unique – there are likely many syndromes (for example, major depressive disorder and essential hypertension) that can arise from many different sequences of hits. These sequences for each syndrome are distinct from each other but partial overlap of sequences may explain disease co-occurrence. Therefore, interventions that decrease hit count would help prevent schizophrenia and multiple other heterogenous syndromes as well.

## Data availability

EEG data and MATLAB code are available from the corresponding author upon request. fMRI data is available from the DecNef Brain Database (https://bicr.atr.jp/decnefpro/).

